# Rationalizing the application of spatial capture-recapture versus space-to-event models for monitoring populations of an endangered social carnivore

**DOI:** 10.64898/2026.09.25.754339

**Authors:** Divyajyoti Ganguly, Avantika Sharma, Uma Ramakrishnan, Arjun Srivathsa

## Abstract

Reliable estimates of population size are fundamental to understanding wildlife population status and informing conservation. Obtaining such estimates remains challenging for unmarked species, particularly in tropical countries where logistical and financial constraints limit intensive monitoring. The endangered dhole (*Cuon alpinus*), a highly social canid without natural markings, is one such species for which robust population estimates remain scarce. We estimated dhole density in the Valparai plateau, a human-dominated agroforest–forest mosaic in India’s Western Ghats. We compared Spatially Explicit Capture–Recapture (SECR) models based on individual identification from fecal DNA and Space-to-Event (STE) models based on camera-trap detections. We additionally applied both approaches to individually identifiable leopards (*Panthera pardus*) to cross-validate STE versus SECR estimates. SECR estimated dhole density at 22 individuals per 100 sq.km, with STE estimates varying substantially depending on how the camera viewshed area was defined. STE estimates based on post hoc viewshed measurements, particularly after reducing the viewshed by 10%, were comparable to SECR estimates. In contrast, STE underestimated leopard density relative to SECR (7 versus 12 per 100 sq.km), although uncertainty intervals overlapped. Ecologically, our results indicate that human-use agroforest mosaics can support dhole densities comparable to, or exceeding, those reported from several Protected Areas, highlighting their potential importance for dhole conservation. More broadly, we argue that STE models may offer a practical and relatively inexpensive approach for baseline monitoring of dholes from camera-trap surveys, but their reliability depends strongly on accurate characterization of the effective viewshed and may vary across species and landscapes.

## 1. Introduction

Assessing the status of wildlife populations is a fundamental building block of conservation research and management. Because wild animals are often elusive and difficult to observe, conventional census methods are rarely effective for obtaining reliable population counts (Mills et al. 2025). Consequently, a range of statistical methods has been developed to estimate wildlife populations with reasonable confidence (Royle et al. 2013). These methods are most robust when applied to ‘marked’ species, wherein individuals can be distinguished based on unique morphological characteristics (Chandler and Royle 2013; Alonso et al. 2015). For ‘unmarked’ species, estimating population sizes remains a longstanding challenge (Chandler and Royle 2013; Gilbert et al. 2021). For species of conservation concern, this knowledge gap has two major implications. First, without reliable estimates of population size, our understanding of fundamental aspects of their biology, like demographic rates and population trajectories, remain limited (Williams et al. 2002). Second, it becomes difficult to design, implement, and evaluate evidence-based conservation interventions, because population responses to such measures cannot be reliably quantified (Karanth et al. 2003). In such cases, wildlife managers instead use less rigorous sources of information, including expert opinions, population indices, and anecdotal guesstimates. These approaches are particularly sensitive to bias and substantially over- or underestimate true population sizes (Sollmann et al. 2013; Mills et al. 2025). Relying on these estimates can compound the conservation challenges faced by threatened species (Karanth 2011). The problem is further amplified when such estimates are used for regional- or continental-scale syntheses which shape global conservation policy (Araújo et al. 2019; Suryawanshi et al. 2019).

Over the last two decades, several approaches have been developed in an attempt to reliably estimate the population sizes of unmarked species. Occupancy-based abundance models estimate abundance proxies by assuming that areas with higher abundance have greater detection probabilities (Royle and Nichols 2003). Count-based N-mixture models use repeated counts at a given site to estimate abundance, assuming that individuals are not double-counted across surveys (Royle 2004). More recently, the Random Encounter Modelling (REM) framework has been the focus of considerable development, primarily because it is less resource intensive and easier to implement. By integrating encounter rates, animal movement speed and the area sampled by a detector (e.g., camera trap), REMs provide a means for estimating population density or abundance without requiring individual identification (Rowcliffe et al. 2008).

However, their underlying assumptions, including random placement of detectors with respect to animal movement and their movement patterns approximating Brownian motion, as in the random movement of gas particles, may constrain the reliability of resulting estimates (Rowcliffe et al. 2008; Pettigrew et al. 2021; Murphy et al. 2024). Space-to-Event and Time-to-Event models are recent extensions of the classical REM framework that reduce the sensitivity of estimates to animal movement rates and incorporate the timing of detections respectively to improve abundance estimation (Moeller et al. 2018). These approaches retain some limitations of classical REMs and require further testing and validation against more robust estimators, such as Spatially Explicit Capture Recapture (SECR) models.

Despite these methodological developments, obtaining field-based estimates of species’ populations in tropical regions remains particularly challenging. Tropical landscapes are often characterized by inaccessible terrain, limited funding, and substantial logistical constraints (Beaudrot et al. 2019; Zwerts et al. 2021). Moreover, many tropical countries in the Global South lack a long history of systematic ecological research and, until recently, have had relatively limited access to requisite expertise, equipment and infrastructure. Consequently, many species remain data deficient, with even robust baseline estimates of population size or density lacking (Martin et al. 2012; Srivathsa et al. 2022). Testing and streamlining newly-developed methods for obtaining baseline population estimates of many under-assessed yet threatened or endangered species in such regions could, therefore, help address a substantial knowledge gap, and offer a foundation for conservation planning and management.

The Asiatic wild dog ‘dhole’ (*Cuon alpinus*) is a highly social canid that lives in packs of 2–25 individuals. Dholes are found across forested landscapes of South and Southeast Asia–– the ‘southern dhole’, with smaller populations distributed in high-altitudes of the Himalayas and across the arid cold-plains of China–– the ‘northern dhole’ (Kamler et al. 2015; Srivathsa et al. 2023). Most populations of the southern dhole are found in protected forests, with trace populations also reported from production agroforests adjacent to forested habitats (Srivathsa et al. 2020a; Tananantayot et al. 2022; Saravanan et al. 2026). Listed as Endangered under the IUCN Red List, expert opinion-based ‘guesstimates’ suggest a global population of ∼10,000 individuals (of which 1000–2000 may be adult, mature individuals capable of breeding; Kamler et al. 2015). Despite their endangered status, focused conservation monitoring or evidence-based management of dhole populations is virtually non-existent. For strategic management and conservation planning, population status is a crucial metric; this aspect of dhole ecology yet remains prominently missing across its range because of three issues–– (i) dholes have no pelage patterns or markings that enable individual identification; (ii) they purportedly occur at low densities through much of their range; common methods applied for multi-species studies are not ideally suited for estimating dhole populations; and (iii) their sociality further complicates the ability to apply population models primarily developed for solitary carnivores (see Emmet et al. 2021).

Camera-trap surveys focused on species like the tiger *Panthera tigris* have routinely produced ‘by-catch’ data on dholes; several studies therefore use these photo-encounters to calculate and present Relative Abundance Index (RAI) as a basic metric of population size (e.g., Charaspet et al. 2019; Widodo et al. 2020). But RAIs can be grossly misleading, and are not conducive for reliable population size-related inferences, especially for making comparisons across space or time (Sollmann et al. 2013; Burton et al. 2015). Dedicated forays into estimating dhole populations are a relatively recent endeavor. In this regard, the earliest explorations involved occupancy-based abundance models (see Selvan et al. 2014) and site-based abundance models (Ngoprasert et al. 2019); these models are rather sensitive to violations of key assumptions––many such survey design assumptions are invariably violated in field-based surveys (Link et al. 2018). The most reliable method, offering the highest flexibility and inferential strength, is identifying individuals from fecal DNA and the application of powerful Spatial Capture–Recapture models (Srivathsa et al. 2021). Despite its utility, this approach remains relatively expensive and requires access to infrastructure and expertise which may not be commonplace across the dhole’s range-countries. Punjabi et al. (2022) undertook a review of population models for dholes and acknowledged all the problems described above; in an attempt to redress the issue, they applied Space-To-Event (STE) and Time-To-Event (TTE) models as potential alternatives (Moeller et al. 2018). While doing so, they report that STE models produced reasonably realistic results, and therefore hold some promise.

Here, we focus on an agroforest plantation–forest mosaic in the Western Ghats landscape of Southern India to estimate dhole populations using two parallel approaches: (1) Spatially Explicit Capture–Recapture models using data from fecal DNA obtained through scat surveys (henceforth, SECR), and (2) STE models using photo-capture data generated through systematic camera-trap surveys (henceforth, STE). Adapting from and building upon the approach by Punjabi et al. (2022), we use the photo-encounter data of leopards *Panthera pardus* obtained from the same camera-trap surveys as ‘control’; we apply both, likelihood-based SECR, and STE models to the leopard data to gauge the reliability of STE models. Finally, we discuss the practicality, logistical feasibility and trade-offs when using SECR-versus STE-based methods for monitoring dhole populations across forest and agroforest landscapes of South and Southeast Asia.

## 2. Materials and methods

### 2.1 Study area

The Valparai plateau, spanning over 220 sq.km in the Anamalai Hills of the Western Ghats in India, is a human-modified tropical landscape and mosaic agroforest system dominated by tea plantations interspersed with rainforest fragments, coffee and cardamom estates, eucalyptus plantations, open grass patches, swamps, streams, and human settlements (Mudappa et al. 2007). Situated at an elevation of 700–1500m above sea level, the plateau receives an average annual rainfall of approximately 2800mm, predominantly during the south-west monsoon, and contains rainforest fragments ranging from 1–300ha embedded within extensive tracts of privately-owned plantations. The plateau has a human population of more than 71,000 people (Census of India 2011, Ministry of Home Affairs) and is surrounded by Protected Areas on all sides: Anamalai Tiger Reserve in Tamil Nadu, Parambikulam Tiger Reserve and Vazhachal Reserved Forest in Kerala, with which it shares contiguous rainforest habitats (Osuri et al. 2019). The plantation–forest mosaic supports a diverse herbivore assemblage, including Asian elephants (*Elephas maximus*), gaur (*Bos gaurus*), sambar deer (*Rusa unicolor*), Indian muntjac (*Muntiacus muntjak*), wild pig (*Sus scrofa*), and mouse deer (*Moschiola indica*), which in turn sustain a carnivore community comprising dholes and leopards, together with sloth bears (*Melursus ursinus*) and a suite of mesocarnivores; tigers are reported occasionally near the boundaries between plantations and the adjacent Protected Areas (Kumara et al. 2004). Dholes are found across the entire landscape, typically residing in the tea plantations and subsisting largely on the abundant wild prey available within the mosaic, the plateau potentially supports 5–6 resident packs, with additional packs or individuals from surrounding forests using its outer edges incidentally (Pious et al. 2025; Saravanan et al. 2026; authors’ personal observations). Leopards are also resident in the landscape, although they likely occur at lower numbers than dholes (Navya et al. 2014).

### 2.2 Field survey design and data collection

Surveys for fecal DNA were conducted during the dry season from December 2023 to March 2024. We surveyed six routes strategically placed across the landscape along foot-trails within tea plantations. The routes, ranging from around 10–17 km each, were divided into 100m segments and surveyed 6–8 times, i.e., temporal replicates (**Fig. 1**). Dhole scats were identified based on size, shape, smell, deposition style, and location (latrine sites; Andheria et al. 2007). Survey teams consisting of 2–3 researchers collected DNA from scats using sterilized swabs. As per protocol, we only collected scats deemed to be ‘fresh’ (classified as such if it was <1 day old), or in some cases when they were classified as ‘between’ (2–3 days old, but the outer layer remained intact because they were deposited under canopy shade). For each piece of scat, we collected samples through two swab-draws (replicates) and stored them in a lysis buffer solution (Longmire et al. 1997; Ramón-Laca et al. 2015). For each scat pile, we recorded GPS coordinates, condition, number of scats in the pile and other descriptive remarks.

**Figure 1.**
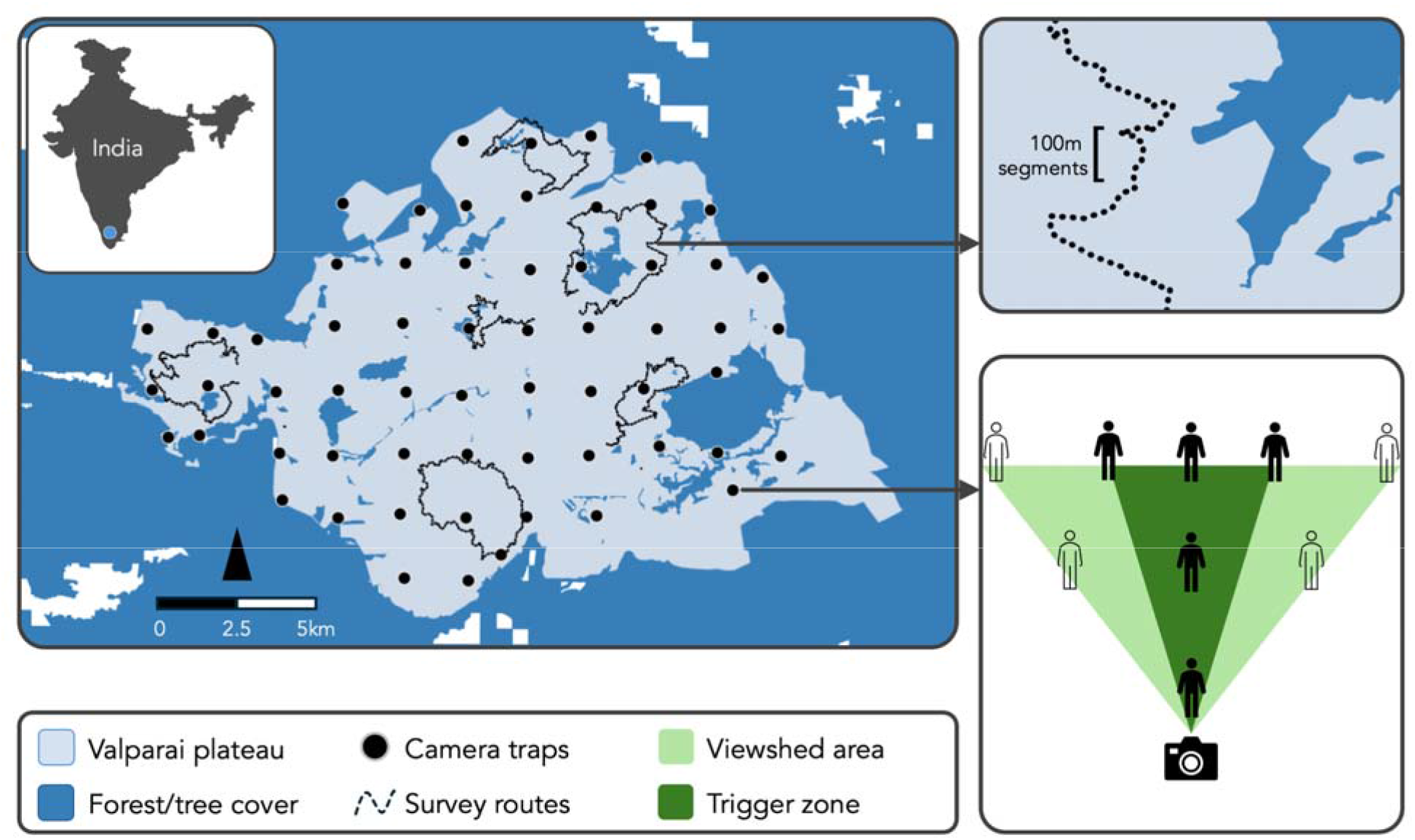
Study area in the Valparai plateau of India’s Western Ghats, a tea-agroforest-dominated landscape, surrounded by forested Protected Areas or reserve forests on all sides. The map shows locations of camera traps and survey routes (inset: location of Valparai in India); Forest/tree cover includes rainforest fragments, shade-grown coffee, eucalyptus and cardamom plantations. Top-right: sampled survey route along a foot-trail, showing 100m segments. Bottom-right: a schematic representation of how the viewshed area was measured; surveyors walked in front of the camera trap(s) to identify the Trigger Zone (solid human figures) and subsequently took more photographs to identify the total viewshed outer limits (outline-only human figures); the viewshed area was then calculated as the outermost bounds of the zone wherein camera-triggers yielded photo-captures of the moving object within the frame.

Systematic camera-trap surveys were carried out from January 2023 to May 2023. For this, we first overlaid a grid network with 61 4-sq.km cells across the Valparai plateau. Cameras were set up within these cells, at or near the centroid, typically along foot-trails in the tea-coffee-cardamom plantations or forest fragments (**Fig. 1**; also see Pious et al. 2025). The specific location was chosen based on our knowledge of dhole behavior and indirect sign-evidence, ensuring that the trap placement would maximize possibilities of dhole detection. In each location, we set up one digital camera (a combination of Cuddeback IR Model H-1453 and Browning Model BTC-7E) at a height of approximately 45cm from the ground. Cameras were active for 11–16 days at a time; each day was treated as a temporal replicate. Given that our resources were limited, we used a block-shifting approach to survey the entire landscape (see Karanth and Nichols 1998) in a short duration, reducing chances of violating closure assumptions needed for unbiased estimates in closed-population SECR models.

### 2.3 Analytical approach for SECR

We leveraged the Single Nucleotide Polymorphism ‘SNP’-based workflow for genotyping and individual identification of dholes from faecal DNA (Srivathsa et al. 2021). This workflow comprises a panel of 150 SNPs. We refined this panel to retain 96 SNP markers that consistently worked across samples. First, DNA was extracted from samples using the Qiagen DNA Extraction kit following a slight modification of the manufacturer’s protocol. Next we undertook multiplex PCR (mPCR) to amplify DNA in the extracts individually using the panel of SNP-specific primers. We then diluted the amplified DNA and subjected it to iPCR where each sample was assigned an unique barcode. The samples were pooled together, purified to remove non-target nucleotide fragments and sequenced using a Illumina miseq kit. We analysed the resultant reads using a standard bioinformatics pipeline: removing barcode sequences (trimming) using TrimGalore! (Krueger, F) with the options: “quality 30”, “phred33”, “stringency 5”, “length 30” and “max_n 1”, mapping the reads onto a reference genome using BWA (Li 2013), sorting the mapped reads using SAMtools (version 1.9; Li et al. 2009) and SNP calling using BCFtools. We used VCFtools to retain SNPs with a minimum site quality of 30, genotype quality ≥30 and depth ≥15 and excluded indels, sites with >50% missing data and sites with a minor allele count <1. Subsequently, samples genotyped at fewer than 30 of the resulting SNPs were excluded from the dataset. Finally, we estimated pair-wise genetic relatedness within and between samples (within– two swab-draws from the same sample; between– each sample paired with every other sample from the entire dataset); the relatedness values range from 0 (samples referring to unrelated individuals) to 1 (samples referring to the same individual), calculated using PLINK (version 1.9; Purcell et al. 2007). Relatedness between the replicate swab-draws for a subset of our samples was used to set a threshold for assigning samples to individual dholes from the entire dataset. Because within-sample relatedness can often be <1 owing to missing or poor quality SNPs, which we also observed in our sample-subset, our threshold was set to ≥0.80 (**Figure S1, Supplementary File 1**). In other words, any two samples whose relatedness was ≥0.80 were deemed to be from the same individual. For leopards, the process was more straightforward; we relied on standard visual identification of individuals from camera trap photo-encounter data based on their unique rosette patterns using software HotSpotter (Crall et al. 2013).

We applied the classical likelihood-based SECR models for data on dholes and leopards to estimate population density (D), baseline capture probability (g0) and movement parameter associated with spatial scale of detection (σ; Borchers and Efford 2008; Efford et al. 2009; Kéry et al. 2010). To do so, we first created a state-space that included a buffer of 10 km from the outermost sampled segments (dholes) or camera-trap locations (leopards). We used the bounds of the state-space to create a grid network of 0.25 sq.km cells. Within this grid network, we retained only those cells that overlapped with putative dhole or leopard habitats, such as forest, agro-forest and plantation; the centroids of these cells collectively defined our habitat mask. Next, we snapped dhole detections along trail segments to the nearest cell centroid; the centroids were now treated as pseudo-detectors (Russell et al. 2012). This enabled us to create a capture history linking dhole captures and recaptures to these centroids. For leopards, we repeated the same process, except that camera locations with captures were snapped to the closest grid-centroid. For both species, we compared the exponential (exp), half-normal (hnm) and hazard half-normal (hhn) models and chose the best-supported one based on AICc scores to derive these parameter estimates. Abundance was estimated within an Effective Sample Area (ESA) by multiplying with the estimated density. The ESA was created by adding a buffer equal to the Estimated σ ✕ √5.99 around the extent of the surveyed routes (Royle et al. 2013; Srivathsa et al. 2015, 2021). Abundance within Valparai was calculated by multiplying the species’ density with the plateau area. All analyses were implemented using the ‘secr’ package in R (R Development Core Team 2026).

### 2.4 Analytical approach for STE

In the STE model, animal detections within the viewshed area of a camera-trap array are used to estimate abundance for a given state-space (Moeller and Lukacs 2022). The STE model is not sensitive to movement rates, but does predicate meeting certain basic assumptions of closed-population models like demographic and geographic closure. Two key prescribed requirements include (i) random placement of camera traps, and (ii) that individual animal observations are independent. Finally, it assumes a 100% probability of photo-capture within the viewshed area; it does not account for a <1 capture probability in its current operational stage. But we note that these models have been applied to studies where surveys were conducted with non-random placement of the camera trap, focusing on territorial species (see Loonam et al. 2021). In fact, Punjabi et al. (2022) showed that despite non-random camera placement, leopard abundance estimated using the STE approach was comparable to the contemporaneous estimate from SECR models. They used this comparison to rationalize the use of the STE model for dholes. We applied this approach to our data to compare densities of both dholes and leopards against their SECR equivalents. The STE approach can be sensitive to the camera’s viewshed area and needs to be tailored to each trap-location (Moeller and Lukacs 2022; Carswell et al. 2025; Koeck et al. 2025). For each camera-trap, we calculated the viewshed area in two ways: (1) based on the trigger distance and width (trigger area), and (2) post hoc measurements informed by the viewable area in camera trap images (see **Fig. 1**). In both the cases, the area is given by the area of a trapezium, where the widths are defined by the trigger or viewable width and the width of the camera lens, and the height is given by the trigger or viewable height. We also included two more variations of the viewable area category by increasing and reducing it by 10%, yielding four types of viewshed areas (Trigger zone, Viewshed area, Viewshed +10%, Viewshed -10%) that were used to estimate dhole and leopard abundance. To run these models we set both the sampling frequency and length to 1 second to ensure instantaneous sampling. STE analyses were implemented using the ‘spaceNtime’ package in R (R Development Core Team 2026); because these models can be computationally intensive, we executed them on a Linux-based institutional cluster.

## 3. Results

A total of 655 camera-trap days of effort yielded 187 dhole and 61 leopard photo-encounters across the Valparai plateau. We identified 10 individual leopards; individuals in some photographs could not be identified because they were blurred or of poor resolution, preventing us from confidently assessing pelage patterns. Of the ten identified individuals, only two were detected across separate occasions and one was detected across two camera-trap stations. The sign surveys for dhole resulted in the collection of 157 fecal samples, of these we retained 121 samples based on SNP quality control filters and identified 61 unique individuals (see **Supplementary File 1** for more details). Of these 61, 21 individuals were detected across multiple occasions and 19 were detected across multiple grid-cells (pseudo-detectors).

While examining models with exp, hnm, and hhm detection functions, the exp model had the lowest AICc score, but differences among models were very small (ΔAICc = 0.00–0.33) and AICc weights were comparable (0.37, 0.31, and 0.31; see **Table 1**). Monte Carlo goodness-of-fit tests indicated no significant lack of fit for any model (exp: *p* = 0.15; hnm: *p* = 0.34; hhn: *p* = 0.34; **Supplementary File 2**). We selected the exp model for the primary analysis based on biological plausibility; estimates from this model were consistent with independent field observations of ∼50 individuals (5–6 packs) in the landscape. Estimates from the alternative detection functions appeared less biologically realistic, despite similar AICc support and adequate goodness of fit. For leopards, the exp model had the best support based on AICc scores and weights (**Table 1**).

**Table 1.** Model selection table of SECR models for dholes and leopards in the Valparai landscape (2023-24). K: number of parameters, Loglik: log likelihood, AICc: Akaike’s Information Criterion corrected for small sample size, ΔAICc: model AICc - minimum AICc, AICcwt: AICc weight.

| Model | K | Loglik | AICc | $\Delta AICc$ | AICcwt |
| --- | --- | --- | --- | --- | --- |
| **Dhole** |  |  |  |  |  |
| exp | 3 | -290.18 | 586.78 | 0.000 | 0.37 |
| hnm | 3 | -290.34 | 587.11 | 0.33 | 0.31 |
| hhn | 3 | -290.34 | 587.11 | 0.33 | 0.31 |
| **Leopard** |  |  |  |  |  |
| exp | 3 | -40.61 | 91.22 | 0.00 | 0.81 |
| hnm | 3 | -42.74 | 95.48 | 4.26 | 0.10 |
| hhn | 3 | -42.79 | 95.58 | 4.36 | 0.10 |

Models: exp— exponential, hnm— half normal, hhn— hazard half-normal

The model-estimated density (D), baseline detection probability (g0) and movement parameter (σ were 21.81 (12.26SE), 0.01 (0.003) and 3757 (2219), respectively, for dholes (**Table 2**; **Figure 2**). For leopards, we estimated D = 11.71 (5.67SE), g0 = 0.34 (0.21) and σ = 364 (93). The abundance of dholes was 46 (range: 20–73) individuals within the Valparai plateau and 161 (range = 70–251) within the ESA (**Table 2**; **Figure 2**). For leopards, the area of the ESA was the same as that of Valparai, yielding an abundance of 25 (range = 13–37) individuals (**Table 2**).

**Figure 2.**
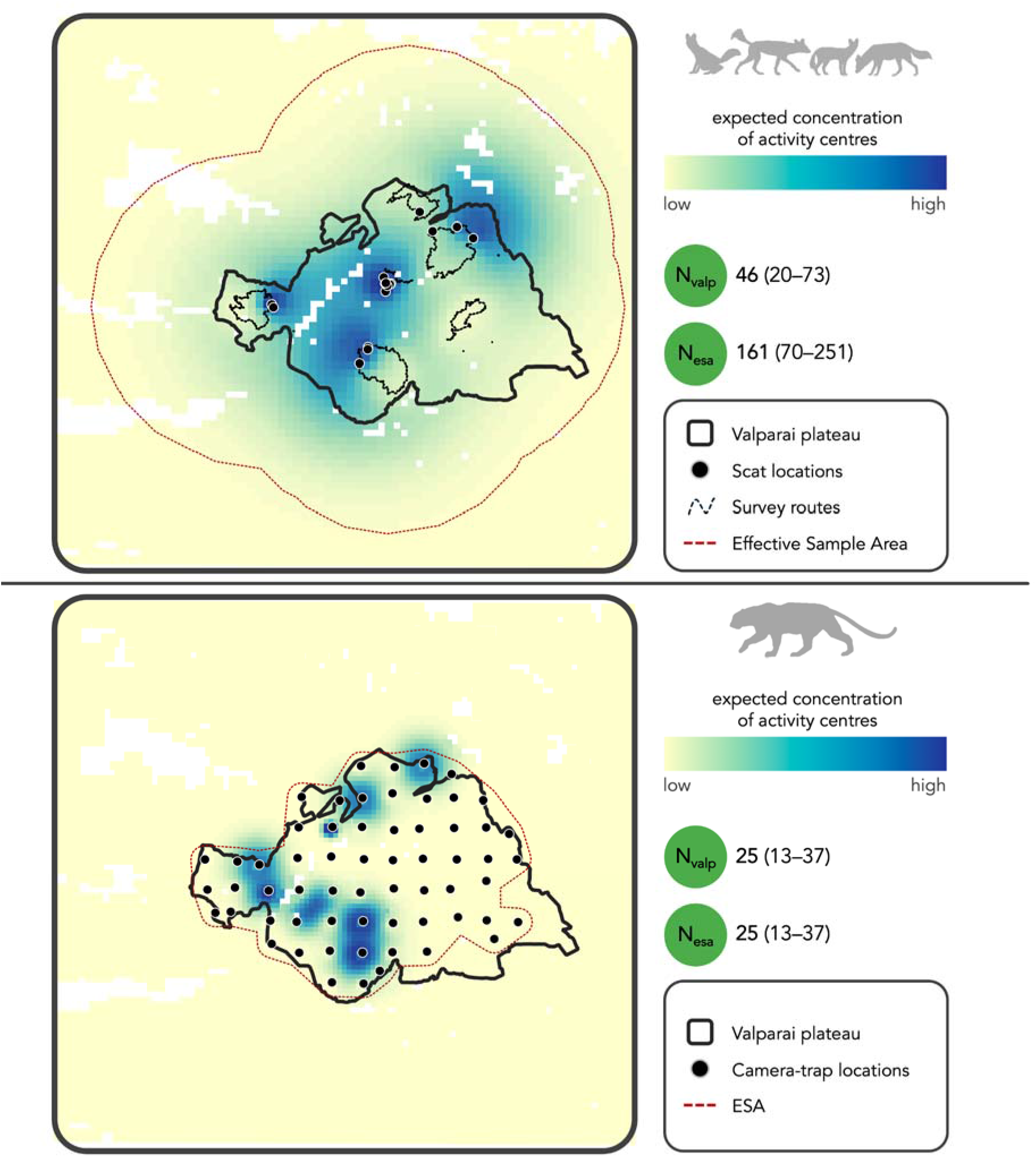
Summed conditional activity-centre probability surface for detected dhole individuals (top panel) and leopard individuals (bottom panel) within the state-space, derived from the fitted SECR models. Darker colours indicate higher expected concentration of activity centres. The dotted line along or beyond the Valparai plateau boundary is the Effective Sample Area (ESA), calculated as √5.99 ✕ distance, applied as a buffer on the outer bounds of the sampled routes. Abundance estimates are presented for the Valparai plateau boundary (N_valp_) and for the ESA (N_esa_).

**Figure 3.**
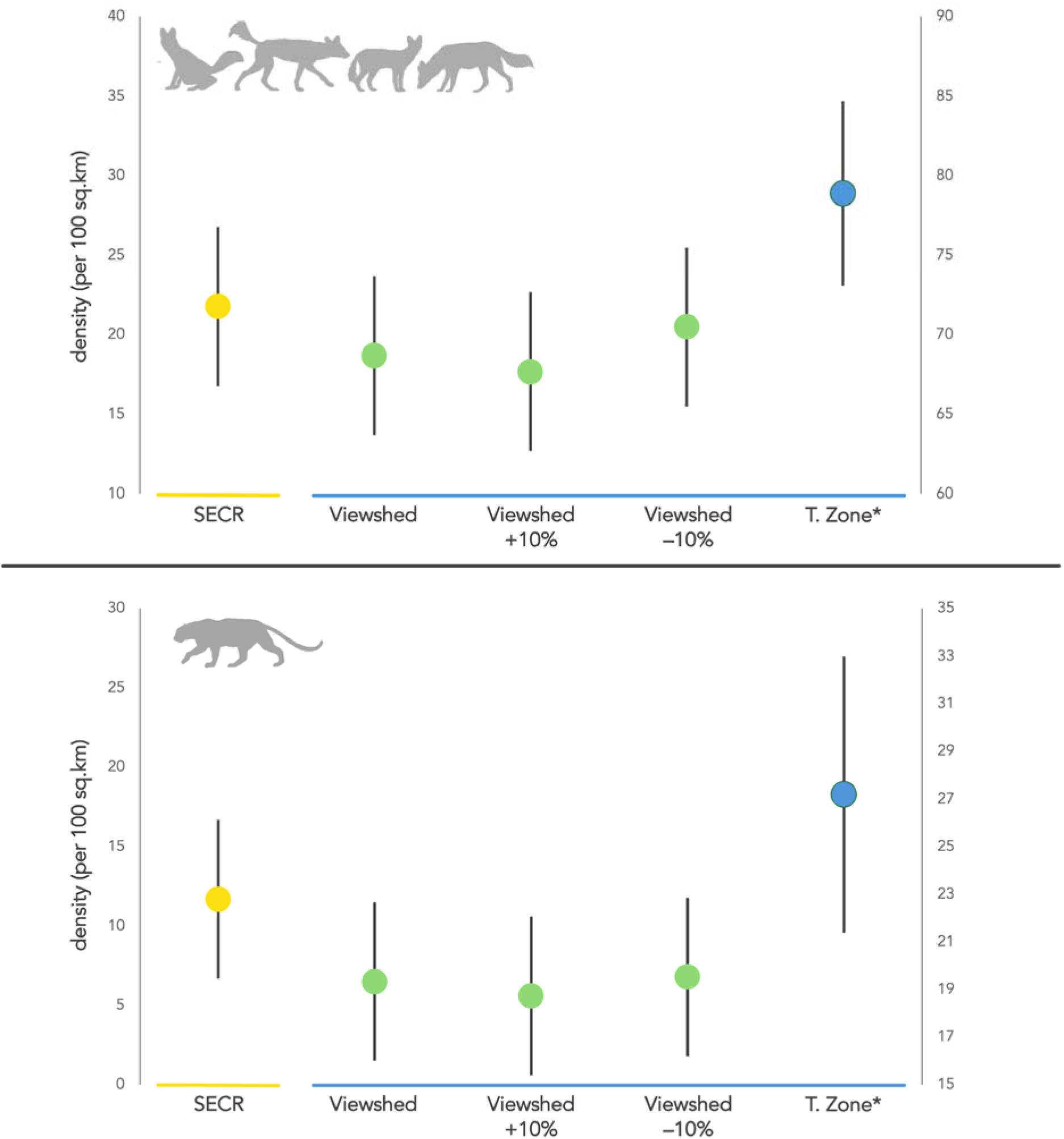
Comparisons of estimated densities (per 100 sq.km; vertical bars are standard errors) for dholes (top panel) and leopards (bottom panel) in the Valparai landscape. For dholes, SECR is from fecal DNA-based identification of individuals; for leopards, SECR estimates are from individual identification through camera-trap photographs; Viewshed, Viewshed+10%, Viewshed–10%, and Trigger Zone (‘T. Zone’) are estimates from variations of viewshed area while using the same STE model; *T. Zone estimates for both species were considerably higher and therefore plotted on the secondary Y-axis.

**Table 2.** Results from SECR models: estimated basal encounter rate (g0), movement parameter (σ) and density (D) for dholes and leopards in the Valparai landscape (2023–24).

| Model | $g_0$<br>(SE) | $\sigma$<br>(SE) | ESA<br>(km<sup>2</sup>) | D (SE)<br>/100km<sup>2</sup> | N<sub>valp</sub><br>(range) | N<sub>esa</sub><br>(range) |
| --- | --- | --- | --- | --- | --- | --- |
| **Dhole** |  |  |  |  |  |  |
| exp | 0.01 (0.003) | 3757 (2219) | 737 | 21.81 (12.26) | 46 (20 – 73) | 161 (70 – 251) |
| hnm | 0.002 (0.001) | 9795 (5054) | – | 12.50 (4.88) | 27 (16 – 37) | – |
| hnn | 0.002 (0.001) | 9794 (5052) | – | 12.50 (4.87) | 27 (16 – 37) | – |
| **Leopard** |  |  |  |  |  |  |
| exp | 0.34 (0.21) | 364 (93) | 213 | 11.71 (5.67) | 25 (13 – 37) | 25 (13 – 37) |
| hnm | 0.12 (0.07) | 654 (125) | 259 | 9.80 (4.26) | 21 (12 – 30) | 25 (14 – 36) |
| hnn | 0.13 (0.08) | 653 (126) | 259 | 9.83 (4.29) | 21 (12 – 30) | 26 (14 – 37) |

The area of the Valparai plateau (valp) used in the abundance calculations is 213 sq.km. Models: exp— exponential, hnm— half normal, hnn— hazard half-normal; Abundance (N) calculated as density × valp area (N<sub>valp</sub>) and density × Effective Sample Area (N<sub>esa</sub>); the min–max range for N was calculated using D–(SE) and D+(SE) and multiplying these with valp area and ESA respectively. Dhole ESAs informed by the $\sigma$ estimates of hnm and hnn models were bigger than the habitat mask, hence, we did not report the ESA and N<sub>esa</sub> estimated by these models.

The STE analyses generated multiple estimates of abundance, which varied based on the viewshed area considered: the Trigger Zone area, viewshed area (measured post hoc) and viewshed area +/–10%. For both dhole and leopard, the abundance (and density) within the Valparai plateau were higher using the Trigger Zone as compared to the other viewshed variations (**Table 3**). Among the post hoc viewsheds, the ‘viewshed+10%’ variant produced a slightly lower dhole abundance estimate than the ‘viewshed’ variant; ‘viewshed–10%’ produced an estimate closest to the SECR estimate (44, range = 41–47; **Table 3**). In the case of leopard, both ‘viewshed’ and ‘viewshed–10%’ variants produced similar abundance estimates, whereas the ‘viewshed+10%’ estimate was slightly lower.

**Table 3.** Results from STE models for dholes and leopards in the Valparai landscape (2023–24). Trigger Zone, Viewshed, Viewshed+10% and Viewshed–10% are estimates from variations of viewshed area calculations.

| Viewshed Type | D (SE)/100km<sup>2</sup> | N<sub>valp</sub> (range) |
| --- | --- | --- |
| **Dhole** |  |  |
| Trigger Zone | 78.87 (5.81) | 168 (156–180) |
| Viewshed | 18.78 (1.37) | 40 (37–43) |
| Viewshed+10% | 17.84 (1.27) | 38 (35–41) |
| Viewshed–10% | 20.66 (1.52) | 44 (41–47) |
| **Leopard** |  |  |
| Trigger Zone | 27.23 (3.40) | 58 (51–65) |
| Viewshed | 6.57 (0.81) | 14 (12–16) |
| Viewshed+10% | 5.63 (0.72) | 12 (10–14) |
| Viewshed–10% | 6.57 (0.87) | 14 (12–16) |

The area of the Valparai plateau (valp) used in the density calculations is 213 sq.km. The min–max range for N<sub>valp</sub> was calculated as N<sub>valp</sub>–(SE) and N<sub>valp</sub>+(SE). Density (D) was calculated by dividing N<sub>valp</sub> by the valp area; SE of D was calculated by dividing SE of N<sub>valp</sub> by valp area.

## 4. Discussion

All estimates of dhole population sizes published hitherto–– the robustness of the method applied notwithstanding–– have been from protected forest habitats. We present perhaps the first such information from a human-use agroforest production area in a globally important conservation landscape for the species. Our results from SECR and STE (especially, field-measured viewshed–10% area) were comparable for dholes, but SECR versus STE differed substantially for leopards. Most previous studies report that higher dhole densities are limited to Protected Areas. On the contrary, our results show that select locations may support exceptionally high densities, and that certain ‘secondary habitats’ like Valparai could behave like population sources rather than as pseudo-sinks for the larger dhole metapopulation in the Western Ghats landscape.

### 4.1 Ecological insights on carnivores in shared spaces

Across landscapes, regions, and spatial scales, dholes have consistently been documented at higher occupancy probabilities (and inferentially, higher densities) within Protected Areas as compared to outside (Karanth et al. 2009; Srivathsa et al. 2014, 2019a, 2019b, 2020a; Thinley et al. 2021; Tananantayot et al. 2022). In this context, our field observations, and subsequently our density estimates from Valparai hint at a certain peculiarity in this location. Dhole presence in commodity agroforest expanses like tea and coffee plantations is fairly well-documented (Gangadharan et al. 2016; Pious et al. 2025; Saravanan et al. 2026). But the densities we report appear comparable or higher than those from other Protected Areas in Southern India, Northeast India and Southeast Asia (Selvan et al. 2014; Ngoprasert et al. 2019; Srivathsa et al. 2021; Punjabi et al. 2022). This apparent anomaly likely stems from a set of local conditions in Valparai that are conducive for high dhole numbers: (1) human–dhole interactions in shared agroforests are extremely benign, as dholes do not attack or harm humans and seldom feed on domestic livestock (Saravanan et al. 2026); (2) Valparai has adequate wild prey densities that sustain larger dhole packs (Kumara et al. 2004; Pious et al. 2025); and (3) the near-absence of tigers and low densities of leopards (Navya et al. 2014; this study) eliminate pressures of top-down competition. Our results pertaining to leopards, on the other hand, closely mirror estimates from a forest–tea-plantation mosaic in Northeast India, where Paul et al. (2024), report somewhat similar densities (5–10 individuals per 100 sq.km). Considered together, our findings underscore the conservation potential of production landscapes that could offer a new genre of biodiversity preservation; these production agroforests represent unique socio-ecological systems that can benefit economic progress while also balancing carnivore conservation in shared spaces.

### 4.2 Reconciling parameter estimates with field realities

We report dhole and leopard abundance at two scales–– Valparai plateau and the Effective Sample Area. Abundance within the plateau can inform ground-level management interventions, while ESA-based abundance refers to the estimated number of activity centres across a spatial extent that is analytically pertinent; the latter figure is useful in cases where there are no clear jurisdiction boundaries defining the study area. Interestingly, our estimate of N_valp_ (∼46) was lower than the number of individuals identified from scats (n = 61). This may be explained by some activity centres projected at or beyond the plateau’s edge, suggesting that dholes from the surrounding Anaimalai and Parambikulam reserves itinerantly use the plateau. The movement parameter (σ) is known to scale inversely with density, whereby individuals are expected to show shorter movement distances (or, smaller home-range sizes) at higher densities (Efford et al. 2016). For Valparai’s dholes, σ was ∼3✕ larger than those reported from Wayanad Sanctuary in the same landscape, while both locations had similar dhole densities (14–19 per 100 sq.km; Srivathsa et al. 2021; A. Srivathsa, unpublished results). Leopards, in contrast, showed extremely limited movement within the plateau; most recaptures were temporal, with only one instance of a spatial recapture (across two locations). We speculate that σ for dholes was biased upwards because of some recaptures across spatially segregated packs/activity centres (up to 13 km apart), indicating frequent wide-ranging movement by a few individuals. This finding corroborates our field observations where we found intensive and frequent deposition of dhole scats at latrine sites along putative edges of pack territories. Such intensive territory markings at road-junctions is characteristic of highly social canids (see Stępniak et al. 2020 for wolves *Canis lupus*; Apps et al. 2022 for African wild dogs *Lycaon pictus*); some lines of evidence from behavior studies imply the same may be the case with dholes (Ghaskadbi et al. 2022). In the absence of high co-predator presence or other sources of top-down control, dholes in Valparai could be actively negotiating space because of higher conspecific competition, resulting in these observed patterns.

### 4.3 The paradox and potential of STE models

Our survey design violated a key assumption of the STE approach–– camera traps were not randomly deployed but placed along trails to maximize carnivore detections. Despite this, dhole density and abundance estimates were comparable to gold-standard SECR estimates. This finding may add to the argument that STE can tolerate some assumption violations without substantially compromising parameter reliability (Loonam et al. 2021; Ausband et al. 2022; Punjabi et al. 2022), but further work is needed to identify thresholds beyond which sensitivity to assumption violations are not tenable. In contrast, STE substantially underestimated leopard density (and implicitly, the abundance), although uncertainty intervals overlapped with SECR estimates. This differs from Punjabi et al. (2022), who reported comparable STE and SECR leopard density estimates in the northern Western Ghats, suggesting that STE performance may vary based on species, locations and underlying true densities. Across both species, Trigger Zone-based STE density was strongly overestimated, whereas the post hoc measurement of viewable area better approximated the camera viewshed. Reducing this area by 10% improved agreement with SECR estimates, highlighting the importance of accurately quantifying the effectively sampled area, as viewshed inaccuracies can bias STE estimates (Lyet et al. 2023; Koeck et al. 2025). Recent REM refinements further address such methodological limitations; Lyet et al. (2023) proposed bootstrap resampling to approximate a time-lapse design from motion-triggered photographs, potentially reducing bias, while Matsuoka et al. (2026) incorporated temporal activity patterns to improve density estimation and assess temporal variation. Future developments could explicitly account for imperfect detection within camera viewsheds (Lyet et al. 2023). Together, these refinements and the relative simplicity of REM-based models could facilitate their wider application to existing large-scale camera-trap datasets, accelerating our ability to make robust status assessments for dholes and similar under-researched species across the tropics.

### 4.4 Practical considerations for generating baseline estimates

Although our estimates of dhole densities from SECR and STE show reasonable agreement, we acknowledge and highlight certain study design caveats which may have influenced the observed patterns. First, our camera-trap surveys and fecal DNA collection surveys were asynchronous. Owing to delays in obtaining research permits, data collection was staggered, with a seven-month gap between the two surveys (camera-trapping was done in January–May 2023; scat collection in December 2023–March 2024). Our comparison is based on a key assumption that dhole population size did not change drastically in the 6–8 month duration. This assumption rests on three factors: (1) from our experience working in the landscape for multiple years, anecdotal evidence hinted at no major changes in dhole numbers in 2023–2024; (2) given adequate prey densities and stable ecological conditions––as is the case in Valparai, wild canid populations can remain remarkably stable across years (e.g., see Davies-Mostert et al. 2015, for African wild dogs; Boyce 2018, for wolves; Srivathsa et al. in review, for dholes); and (3) the near-absence of tigers, low densities of leopards (this study) and low conflict with humans (Saravanan et al. 2026) eliminate all potential top-down controls for dhole populations in the landscape. Second, while our overall results reiterate the utility of using STE models for estimating dhole populations, we also note that our field site and results contextually mirror the study location of Punjabi et al. (2022), in that the underlying dhole densities and pack sizes are at the higher end of the scale. We argue therefore that the STE model needs to be thoroughly evaluated in locations with much lower dhole densities, and places where pack sizes are considerably smaller (e.g., Northeast India and large parts of Southeast Asia; Srivathsa et al. 2020b, 2023; Bhandari et al. 2021) to ensure the model’s wider applicability as a standard protocol for conservation monitoring of dhole populations.

## 5. Conclusion

From the perspective of conservation status assessment, our results add an important data point on dhole density from a non-protected area, in particular, a human-dominated agroforest mosaic. Given the paucity of dhole population estimates globally, this holds implicit merit, while also highlighting the incredible potential of secondary habitats in certain locations to sustain viable dhole populations. We nonetheless urge caution and the need to be conservative when interpreting true abundance from STE models; considering the wide variation in numbers for plateau limits versus the Effective Sample Area, using ‘density’ as the primary metric may have more utility for multi-year or multi-location assessments. A worthwhile exercise in this regard would also be to implement STE models in captivity or controlled exclosures, where the number of individuals is known and viewshed area measurements can be quantitatively calibrated, instead of relying on the post hoc adjustments that we show here (with percentage increases– decreases as plausible variations). More broadly, our results have implications for initiating first-time assessments and for monitoring populations in several parts of the ‘southern dhole’ distribution range. We believe the approach described here can be integrated into the IUCN Red List assessment and global guidelines for establishing baseline dhole populations. Finally, given the widespread use of trail-based camera trap surveys across large parts of South–Southeast Asia, other plausible species for which the STE approach may be useful need to be explored, while also ensuring that it is adequately calibrated with reliable, robust methods, as demonstrated in this study.

## Acknowledgements

We express gratitude to the Tamil Nadu State Forest Department for providing research permits [Proc. Nos. WL5(A)/2653/2023 and 878/2023/F1; Permission No: 17/2023] and for their support in implementing the study. We thank M. Shahir, M.A. Shaikh, A. Pious, and A. Das for assistance with field surveys, and M. Puri for generously donating camera traps and other field equipment. We are grateful to S. Darshan, P. Praveen, I. Sinha, A. Chandramouli, V. Paynter, J. Manazhi, L.W. Havmøller and D. Bajaj for all their help in procuring samples, generating data, and conducting laboratory work and analyses. We thank A. Pandit, C. P. Lakshminarayanan and S. Raj from the Next Generation Genomics Sequencing Facility, Bangalore Life Science Cluster (BLiSC) for the Illumina sequencing runs, the NCBS data cluster (supported under project no. 12-R&D-TFR-5.04-0900 by the Department of Atomic Energy, Government of India), and P. Dey from the NCBS Collections Facility and NCBS Wildlife Program for providing working space. We are grateful to Wildlife Conservation Society–India, National Centre for Biological Sciences–TIFR (facilitated by U.R.) and the Rufford Foundation for funding the study and/or providing institutional support. A.S. was supported by the Department of Science and Technology–Government of India’s Innovation in Science Pursuit for Inspired Research (INSPIRE) Faculty Award.

## Supplementary Files

**Supplementary File 1.** Summary of samples collected, the number of SNPs, genotyping success (%), unique individuals identified; distribution of pair-wise genetic relatedness among all sample pairs across the Valparai plateau from 2023 to 2024.

**Supplementary File 2.** Details of Monte Carlo goodness-of-fit test conducted to compare fit of dhole SECR models with exponential (exp), half-normal (hnm) and hazard half-normal (hnn) detection functions.

## Notes

### Competing Interest Statement

The authors have declared no competing interest.

