## Supplementary File 1 for "Rationalizing the application of spatial capture-recapture versus space-to-event models for monitoring populations of an endangered social carnivore"

**Supplementary File 1: Summary of samples collected, the number of SNPs, genotyping success (%), unique individuals identified; distribution of pair-wise genetic relatedness among all sample pairs across the Valparai plateau from 2023 to 2024.**

During the survey period (November 2023–March 2024), we collected 157 dhole faecal DNA samples from the Valparai plateau. SNP calling yielded 60 out of the 96 SNPs across the samples. Of the total samples collected, 121 samples were retained after filtering for quality control, resulting in 77% genotyping success. Based on the 121 samples, we identified 61 dhole individuals.

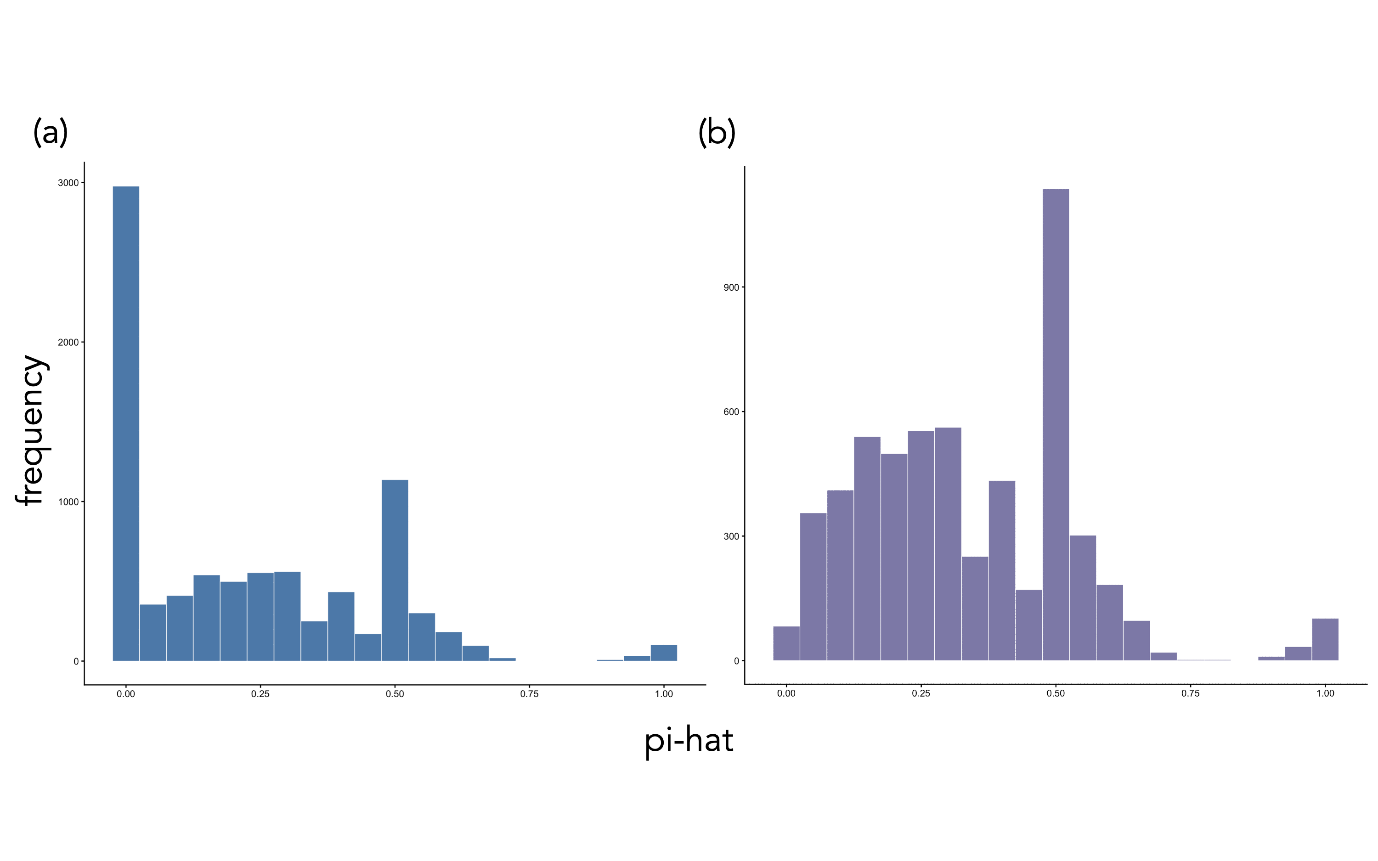
**Figure S1**: Distribution of pair-wise genetic relatedness (pi-hat) among all sample pairs collected in the Valparai plateau (2023–24). (a) distribution of all relatedness scores across sample pairs including ‘zeroes’ (i.e., completely unrelated individuals); (b) distribution of all relatedness scores across sample pairs excluding ‘zeroes’, plotted for easier visualization and interpretation of the non-zero scores (with peaks at 0.25–half-sibs, 0.5–sibs, and 1–recaptures of the same individual).
