## Supplementary File 2 for "Rationalizing the application of spatial capture-recapture versus space-to-event models for monitoring populations of an endangered social carnivore"

**Supplementary File 2: Details of Monte Carlo goodness-of-fit test conducted to compare fit of dhole SECR models with exponential (exp), half-normal (hnm) and hazard half-normal (hnn) detection functions.**

To assess the goodness of fit of our exponential (exp), half-normal (hnm), and hazard half-normal (hhn) detection-function models, we conducted a Monte Carlo goodness-of-fit test using the secr.test() function in the secr package in R (Efford 2011). In this approach, the *f1* statistic, representing the proportion of individuals detected on only one occasion, is compared with its null distribution generated by simulating datasets under each fitted model. The *p*-value represents the rank of the observed statistic within the distribution of simulated values; *p* <0.05 indicates that the observed value is significantly unlikely under the fitted model, providing evidence for the lack of model fit.

Across 999 simulations, none of our three models showed evidence of significant lack of fit based on the *p*-values of the *f1* statistic (Table S1).

**Table S1:** Summary of Monte Carlo goodness-of-fit results for the dhole SECR models. AICc: Akaike’s information criterion adjusted for small sample size; *p*: *p*-value of the *f1* statistic.

| **Model** | **Log likelihood** | **AICc** | ***p*** | **Inference** |
| --- | --- | --- | --- | --- |
| exp | -290.18 | 586.78 | 0.15 | no evidence  for lack of fit |
| hnm | -290.34 | 587.11 | 0.34 | no evidence  for lack of fit |
| hhn | -290.34 | 587.11 | 0.34 | no evidence  for lack of fit |
| Models: exp–– exponential, hnm–– half normal, hhn–– hazard half-normal | | | | |
